# Apoptotic endometrial caspase-3 mediated phospholipase a2 activation, a critical component in programing uterine receptivity

**DOI:** 10.1101/2020.06.29.177337

**Authors:** Sicily E. Garvin, Chandrashekara Kyathanahalli, Arren E. Simpson, Jennifer C. Condon, Pancharatnam Jeyasuria

**Affiliations:** Division of Reproductive Endocrinology and Infertility, Wayne State University; Department of Obstetrics and Gynecology, School of Medicine, Wayne State University, Detroit, Michigan, USA; Department of Physiology, School of Medicine, Wayne State University, Detroit, Michigan, USA

**Keywords:** Apoptotic Caspase-3, endometrium, prostaglandins, luteolysis, uterine receptivity

## Abstract

The objective of this study was to determine the consequence of uterine apoptotic caspase-3 activation on day 1 post coitus (dpc) in the pregnant mouse. We previously demonstrated that during pregnancy uterine caspase-3 activation isolated to the myometrial compartment is largely non-apoptotic and controls uterine quiescence. In this study we determined that uterine caspase-3 activation on 1 dpc may play a critical role in regulating endometrial PGE2 synthesis though iPLA2 activation. These analyses provide novel insight into the molecular mechanisms that regulate previously reported increases in endometrial PGE2 synthesis in very early pregnancy, that act to enhance uterine receptivity.

We have identified the site and impact of that uterine apoptotic caspase-3 activation utilizing uteri isolated from non-pregnant control animals at estrous and diestrous and from control pregnant mice at 1-19 dpc. In addition, uteri were isolated from non-ligated controls (GD), unilateral (UL) and bilateral ligated (BL) uterine horn mouse models at 1, 3 and 6 dpc. Uteri were examined for apoptotic indices, such as caspase-3 activation and TUNEL staining. Immunohistochemical analysis was performed to identify the site of apoptotic caspase-3 activation. The presence of the truncated form of phospholipase A2 (tiPLA2) was examined as a measure of apoptotic caspase-3 mediated iPLA2 activation.

Our analysis determined that apoptotic caspase-3 and iPLA2 activation were limited to the endometrial compartments of the control and unilateral uteri on 1dpc and were not found in the bilateral ligated uterine horn on 3 or 6 dpc. Our data indicates that the presence of a conceptus on 1 dpc triggers an increase in endometrial apoptotic caspase-3 mediated iPLA2 activation. iPLA2 when activated causes the hydrolysis of fatty acids resulting in arachidonic acid release and production of PGE2, which in early pregnancy has been demonstrated to act in a leutoprotective manner, prolonging progesterone synthesis and promoting uterine receptivity.

## Introduction

Despite many advances in assisted reproductive technologies (ART), implantation rates continue to remain low. We know that the process of implantation requires a reciprocal interaction between blastocyst and endometrium, involving both the embryo, with its intrinsic molecular program of cell growth and differentiation, and the progressive differentiation of endometrial cells to accomplish a state of appropriate uterine receptivity. However, it remains unresolved why implantation failure occurs. Failure to implant is thought to occur as a consequence of impaired embryo developmental potential, deficiency in uterine receptivity and/or a diminished embryo–uterine dialogue. Therefore, gaining a better understanding of the molecular events that govern uterine receptivity and implanation is warranted. By understanding the activity and function of the factors involved, it may be possible to use them as predictors of endometrial receptivity or embryo quality to maximize implantation rates in future ART cycles.

Previous analysis has demonstrated in several animal models that an increase in uterine prostaglandin PGE2 biosynthesis in very early pregnancy plays a role in the maintenance of the corpus luteum and luteal function [1]. Uterine PGE2 biosynthesis in very early pregnancy has been demonstrated conclusively to increase ovarian progesterone synthesis through binding its EP2 receptors on the corpus luteum [2, 3]. These events have been demonstrated to increase ovarian progesterone production in early pregnancy, promoting uterine receptivity and implantation. However, the trigger for this early uterine prostaglandin biosynthesis prior to implantation remains unknown.

Our current analysis suggests that a spike in apoptotic caspase-3 activity isolated to the endometrial compartment on 1dpc may trigger this leutoprotective, prostaglandin dependent processes through truncation and activation of iPLA2. Our previous analysis has demonstrated abundant caspase-3 activity in the pregnant uterus across gestation, however its action remains strictly non-apoptotic, tocolytic and is limited to the myometrial compartment [4, 5]. We have also demonstrated a surge in endometrial apoptotic caspase-3, at term, triggers the activation and truncation of uterine iPLA2 and consequently the upregulation of uterine endometrial prostaglandin synthesis [6]. In this current study we propose that the presence of a conceptus triggers a preimplantation spike in endometrial caspase-3 activity at 1 dpc. Which through truncation and activation of iPLA2 may play a critical and central role in leutoprotective prostaglandin synthesis observed in early gestation in multiple species.

## Materials and Methods

### Pregnant Control, Unilateral and Non Pregnant Bilateral Mouse Models

All animal procedures were performed with the ethical and technical approval of the Institutional Animal Care and Use Committee (IACUC) of Wayne State University, protocol number #17-06-285 and in accordance with the Eighth Edition of the Guide for the Care and Use of Laboratory Animals. All lab members involved in animal research received specialized training from Laboratory Animals Medical Services, the centralized animal care program at Wayne State University. Timed pregnant mice (6-8 week old) were purchased from Charles River Laboratories (Wilmington, MA) and maintained with standard pellet diet and water *ad libitum*. All mice were kept in the same room, with a 12/12 hour light/dark cycle.

Under general anesthesia, virgin female ICR mice (4-6 weeks old) (obtained from Charles River Laboratories) through a flank incision, underwent either unilateral or bilateral tubal ligation, proximal to the oviduct to ensure that they subsequently became pregnant in only one horn or could not become pregnant, respectively [7]. Animals were allowed to recover from surgery for at least 7 days before mating. The following criteria were utilized as indications for immediate euthanasia: respiratory distress/failure, aberrant inactivity with obvious discomfort, severe apparent pain and/or recommendations from vivarium veterinarian staff. If any of these criteria were met, the animal would be immediately euthanized. Euthanasia would be performed using medical-grade compressed CO_2_. The displacement rate was 10–20% of the chamber cage volume/minute regulated by a flow meter. Consistent with the above guidelines, housing and nesting material were included in cages, and any apparent discomfort was immediately reported with appropriate intervention. Animals were observed by a lab member at least three times every 24 hours and by an animal technician at least once every 12 hours. In total, 0 animals were euthanized due to respiratory distress/failure, aberrant inactivity with obvious discomfort, severe apparent pain and/or recommendations from vivarium veterinarian staff.

Uterine samples from pregnant control ICR mice (6-8 week old) obtained from Charles River Laboratories were sacrificed 1, 2 3, 4, 5, 6, 8, 10, 11, 13, 14, 15, 17, 18 and 19 dpc (n=3 for each gestational time point) between 800-1000am. Uterine samples were also isolated at estrous, and diestrous from non-pregnant ICR control mice, and from the gravid and non-gravid horns of the unilateral (n=3) at 1, 3 and 6 dpc and bilateral (n=6) mouse models at 1 dpc. For all pregnant and non-pregnant time points examined, uterine tissue was collected from three different animals (n=3) and analyzed separately. All gravid uterine horns were cleared of embryonic material and washed in 1XPBS. Uteri were either flash frozen in liquid nitrogen for subsequent protein analysis, immunofluorescence and TUNEL studies, or fixed in 4% PFA overnight and subsequently embedded in paraffin blocks for sectioning. The Institutional Animal Care and Use Committee of the Wayne State University Detroit approved all animal studies. Euthanasia was performed on all mice at the experimental endpoints described above, using medical-grade compressed CO_2_. The displacement rate was 10–20% of the chamber cage volume/minute regulated by a flow meter. Consistent with the above guidelines, housing and nesting material were included in cages.

### Mouse Estrous Tracking

Estrous and diestrous were identified by a vaginal swab followed by vaginal cytology for a minimum of 2 weeks before the animals were sacrificed. Briefly the vaginas of the mice were washed with 100μl of PBS and isolated cells were analyzed under a dissecting microscope. The stage in the estrous cycle was determined by the proportion of cornified epithelial cells, nucleated epithelial cells, and leukocytes [8]. As the stage of the cycle advances to estrous, mostly cornified epithelial cells are present. Diestrous is the longest of the stages lasting more than 2 days. Vaginal swabs during diestrous show primarily polymorph nuclear leukocytes and a few epithelial cells during late diestrous.

### Subcellular fractionation

Cytoplasmic and nuclear protein extracts were prepared from frozen mouse uterine tissues. In brief, uterine tissue was pulverized in liquid nitrogen and homogenized (IKA homogenizer) in ice-cold NE1 buffer [10 mM Hepes pH 7.5, 10 mM MgCl2, 5 mM KCl, 0.1% Triton X-100 and protease/phosphatase inhibitor mixture (#88669; ThermoFisher scientific)]. The homogenate was centrifuged at 3000 × g for 6 min at 4 °C, and the supernatant was retained as the cytoplasmic fraction. The pellet was washed with NE1 buffer and resuspended in ice-cold NE2 buffer (25% glycerol, 20 mM Hepes pH 7.9, 500 mM NaCl, 1.5 mM MgCl2, 0.2 mM EDTA pH 8.0 containing protease/phosphatase inhibitor). The samples were incubated at 4°C in an Eppendorf thermomixer with vigorous shaking (15 sec/1400 rpm every 5 min) for 1 hr and then centrifuged at 10,600 × g for 10 min at 4°C. The supernatant was retained as the nuclear protein extract.

### Western blot analysis

Equivalent amounts of protein determined by Bicinchoninic acid protein assay kit were resolved by NuPAGE 4–12% Bis-Tris gel (ThermoFisher scientific) electrophoresis and blotted to Hybond-P PVDF membranes (GE Healthcare Bio-Sciences, Pittsburgh, PA, USA). Blots were blocked with 5% nonfat dry milk in Tris-saline buffer (pH 7.4) containing 0.1% Tween-20, and then probed with the following primary antibodies: anti-Cleaved Caspase-3 1:250 (#9664 from Cell Signaling Technology and anti-iPLA2 1:500 (Cat 07-169-i) from Millipore. Immunoreactivity was detected using HRP-conjugated secondary antibody (1:5000), and the bands were visualized using an ECL detection system (ThermoFisher Scientific). The membranes were probed with anti-NCOA3 (for nuclear protein; 1:5000, #PA1-845; ThermoFisher Scientific) or anti-PDI (for cytoplasmic protein; 1:1000, #3501 Cell Signaling Technology, Danvers, MA, USA) to quantify the relative protein expression level. Images of the bands were scanned and analyzed using ImageJ software for relative optical density (ROD) (NIH, Bethesda, MD, USA).

### Immunohistochemistry and immunofluorescence

Mouse uterine tissue was fixed in 4% paraformaldehyde, embedded in paraffin, and sectioned at 5-μm sections and collected on superfrost slides (Fisher Scientific). Parafiin slide sections were deparaffinized and rehydrated through a xylene/alcohol series followed by antigen retrieval by pressure cooking on full power for 21 min in 10 mM citrate buffer (pH 6.0). Tissue sections were blocked in 5% normal donkey serum and incubated in primary antibody anti-cleaved CASP3 (1:100 dilution, #9664 Cell Signaling Technology, Danvers, MA, USA) overnight at 4°C. Sections were washed and incubated with goat anti-rabbit biotinylated secondary antibody (1:1000) using the Vectastain ABC Elite Kit (Peroxidase Goat IgG). Sections were stained using the AEC HRP Detection Kit (Vector Laboratories, Burlingame, CA) following the manufacturers protocol. Nuclei were counterstained blue with haemotoxylin for 2 minutes.

### TUNEL Staining

Paraffin embedded uterine tissues were sectioned at 5μm intervals and collected on Superfrost Plus slides (Fisher Scientific, Pittsburgh PA). Paraffin sections were deparaffinized through xylene and rehydrated through an alcohol series. Sections were permeabilized with 20ug/ml Proteinase K for 8 – 10 mins at room temperature and were stained with the DeadEnd Fluorometric TUNEL assay system as per manufacturers instructions (Promega, #G3250, Madison, WI, USA).

### Statistical Analysis

All data are representative of at least three individual experiments performed in triplicate. Immunoblots were analyzed by densitometry using ImageJ (NIH, Bethesda MD). Statistical analysis of immunoblots was performed with StatPlus:mac software 2009 version (AnalystSoft Inc.). Mouse gestational data points were subjected to a one-way ANOVA followed by pairwise comparison (Student-Neuman-Keuls method) to determine differences between groups. *p* values <0.05 were considered significant.

## Results

### Distinct profile of uterine caspase-3 activation during mouse pregnancy

Western blot analysis was performed on cytoplasmic extracts isolated from uterine tissues originating from non-pregnant mice at diestrous and pregnant mice from 1 to term (19 dpc) (n=3 for each gestational time point (A, B and C)). A significant increase in cleaved caspase-3 on 1 dpc (20 fold) was observed in comparison to diestrous as indicated by the appearance of 19 and 17 kDa fragments. Post 1dpc cleaved caspase-3 levels decline and remain at almost undetectable levels until a mid gestational surge observed between 5 and17 dpc, after which it declines again towards term (Figure 1). PDI is utilized as a loading control to confirm equal loading of cytoplasmic proteins.

**Figure 1.**
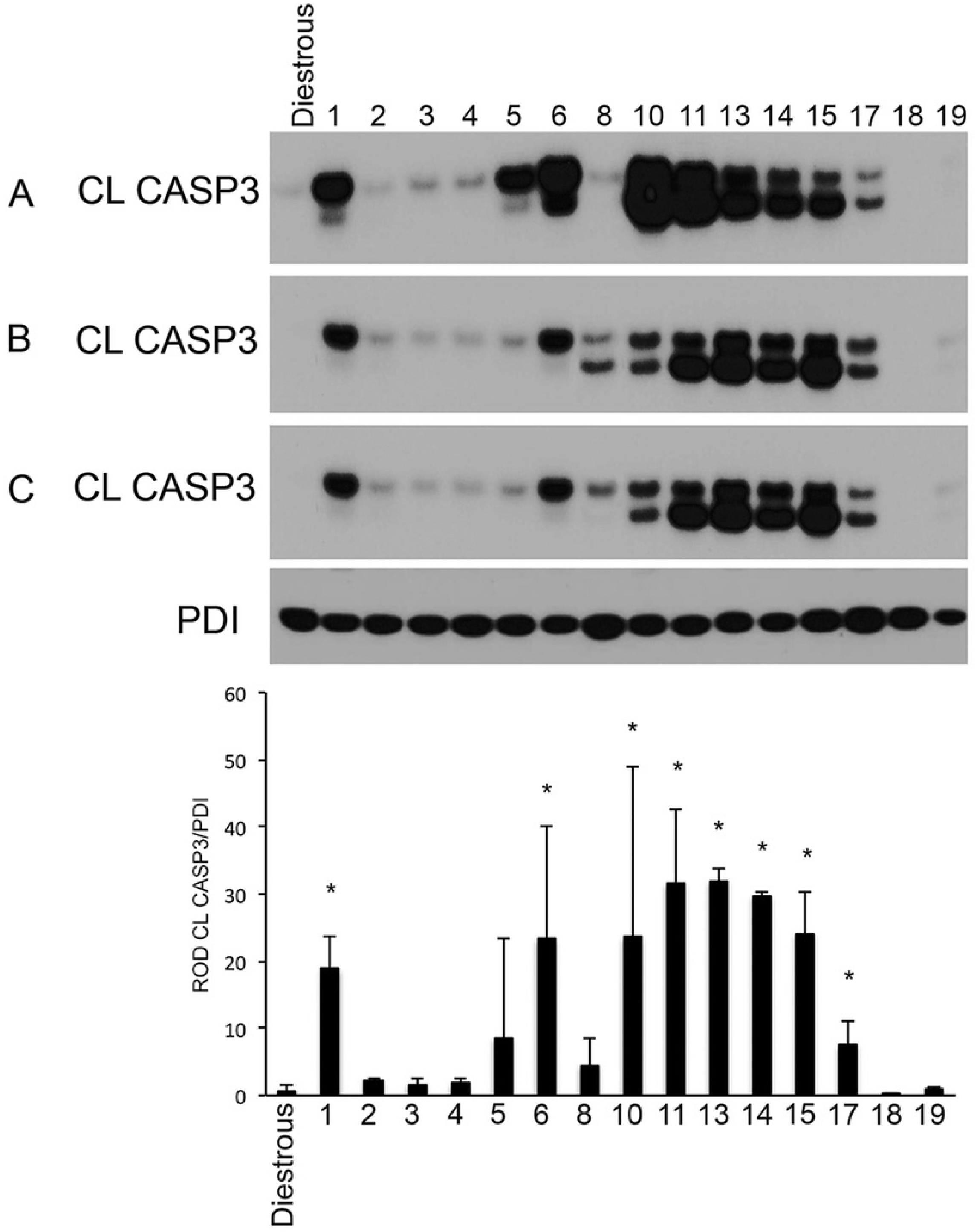
A gestational profile of uterine cleaved caspase-3 (CL CASP3) in the pregnant mouse. Western blot analysis for CL CASP (17/19kDa) in mouse uterine tissue collected at diestrous and from 1-19 days post coitus (dpc) (n=3 for each gestational time point). Protein disulfide isomerase (PDI) is utilized as a cytoplasmic protein loading control. Representative blots from this experiment are shown. Statistical comparisons were performed using one-way ANOVA, and subsequent Newman–Keuls multiple-comparison tests. Data labeled with * are significantly elevated in comparison to diestrous (*p* < 0.005).

### Caspase-3 activation in early pregnancy is apoptotic in nature

Apoptotic caspase-3 activation as identified by the presence of CL CASP3 in the cytoplasmic fraction and cleaved PARP (CL PARP) in the nuclear fraction was examined in mouse uterine tissues isolated from our bilateral ligated mouse model (BL) at 1 dpc, control non-ligated mice (GD) at estrous, diestrous at 1, 3 and 6 dpc and our unilateral ligated mouse model (UL) gravid (P) and non-gravid (NP) uterine horns at 1, 3 and 6 dpc (Figure 2). A significant surge in apoptotic caspase-3 activation as indicated by the appearance of CL PARP is largely isolated to the control GD and the UL, P and NP uterine horns at 1 dpc. Apoptotic caspase-3 activation was greatly reduced in the BL, estrous and diestrous non-pregnant controls and at 3 and 6 dpc in the GD and UL uterine horns, as indicated by the reduced levels of CL PARP. The minor levels of apoptotic caspase-3 activation observed in the BL uterine horns at 1 dpc, strongly confirms that coitus is not the trigger for the increased uterine apoptotic caspase-3 activity and that the surge in uterine apoptotic caspase-3 activation at 1 dpc observed in our GD and UL uterine horns occurs due to the presence of a conceptus. These data also suggest that systemic factors secreted by the conceptus on 1dpc trigger apoptotic caspase-3 activation as CL CASP3 and CL PARP are observed in both the gravid and non-gravid uterine horns of the UL pregnant model. PDI and NCOA3 are utilized as our loading controls for our cytoplasmic and nuclear proteins respectively.

**Figure 2.**
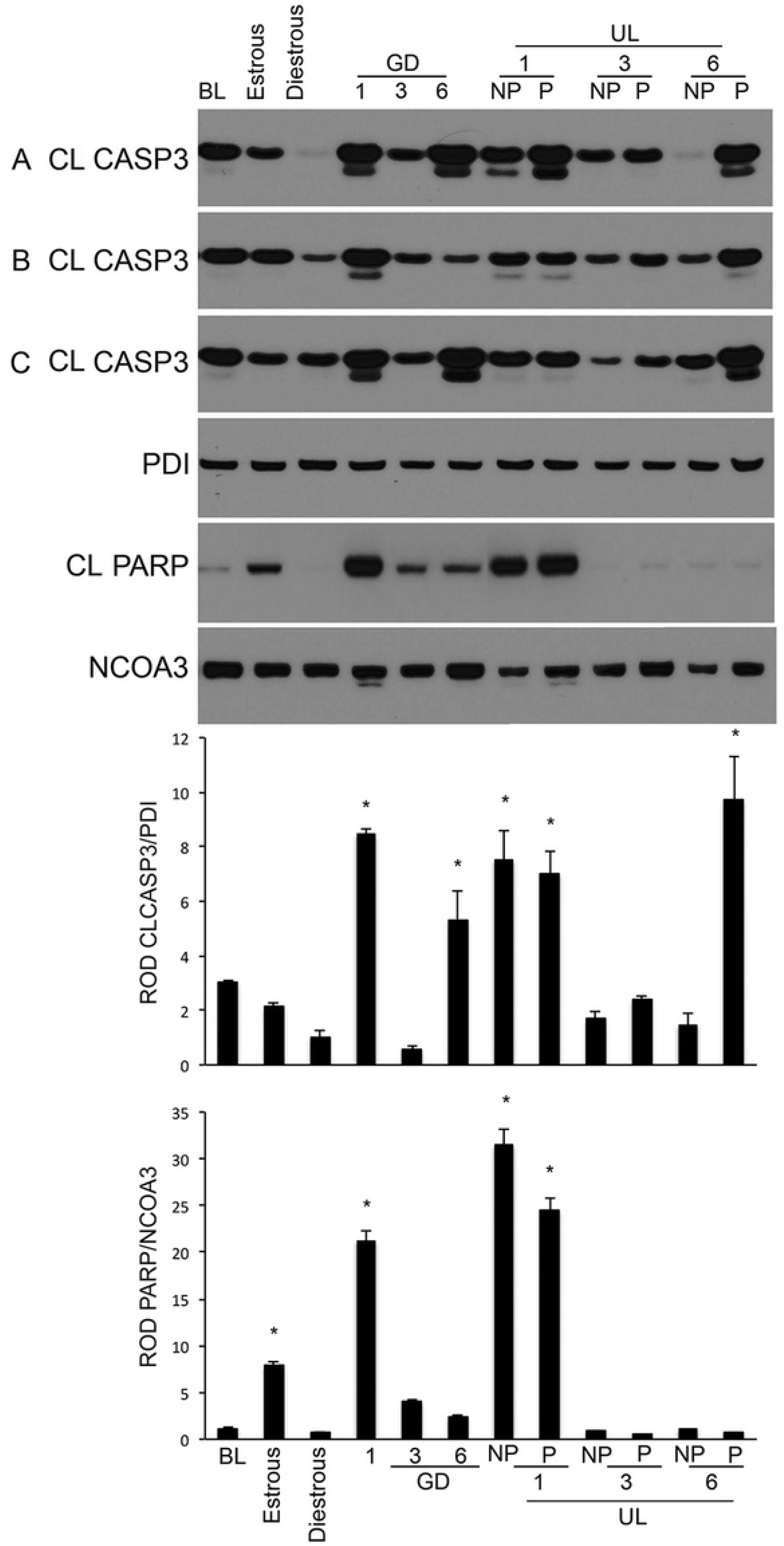
Uterine cleaved caspase-3 (CL CASP3) and cleaved poly ADP-ribose polymerase (CL PARP) levels in control, unilateral and bilateral pregnant mouse models. Western blot analysis of cleaved caspase-3 found at 17/19 kDa and cleaved PARP found at 89kDa in mouse uterine tissue collected from bilateral ligated (BL) at 1 dpc, estrous, diestrous, and control pregnant mice at 1, 3, 6 dpc and unilateral ligated (UL), gravid and non gravid horns (P and NP respectively) on 1, 3, 6 dpc (n=3 for each gestational time point). Protein disulfide isomerase (PDI) is utilized as a cytoplasmic protein loading control and nuclear receptor coactivator 3 (NCOA3) is utilized as a nuclear protein loading control. Representative blots from this experiment are shown. Statistical comparisons were performed using one-way ANOVA, and subsequent Newman–Keuls multiple-comparison tests. Data labeled with * are significantly elevated in comparison to BL uterine horn samples (*p* < 0.005).

### Apoptotic caspase-3 triggers activation of uterine phospholipase A2

The appearance of truncated iPLA2 (tiPLA2) as a measure of apoptotic caspase-3 mediated iPLA2 activation [6] was investigated by western blot analysis of mouse uterine tissues isolated from our bilateral ligated mouse model (BL) 1dpc, control non-ligated mice (GD) at estrous, diestrous, 1, 3 and 6 dpc and our unilateral ligated mouse model (UL) gravid (P) and non-gravid (NP) uterine horns at 1, 3 and 6 dpc (Figure 3). As can be observed apoptotic caspase-3 activation was strongly associated with significantly elevated levels of tiPLA2 on 1dpc in the GD and UL (P and NP) uterine horns. Non-apoptotic caspase-3 activation as observed in Figure 2 (independent of PARP cleavage) did not have the capacity to cleave and activate iPLA2 (Figure 3) as is evident at 6 dpc in the both the GD and UL uterine horns. These data demonstrate in vivo that activation of uterine apoptotic caspase-3 and the cleavage and activation of uterine iPLA2 on 1 dpc occur in tandem and does not occur in the presence of non-apoptotic caspase-3. PDI and NCOA3 are utilized as our loading controls for our cytoplasmic and nuclear proteins respectively. Estrous samples are utilized as our positive control [9, 10].

**Figure 3.**
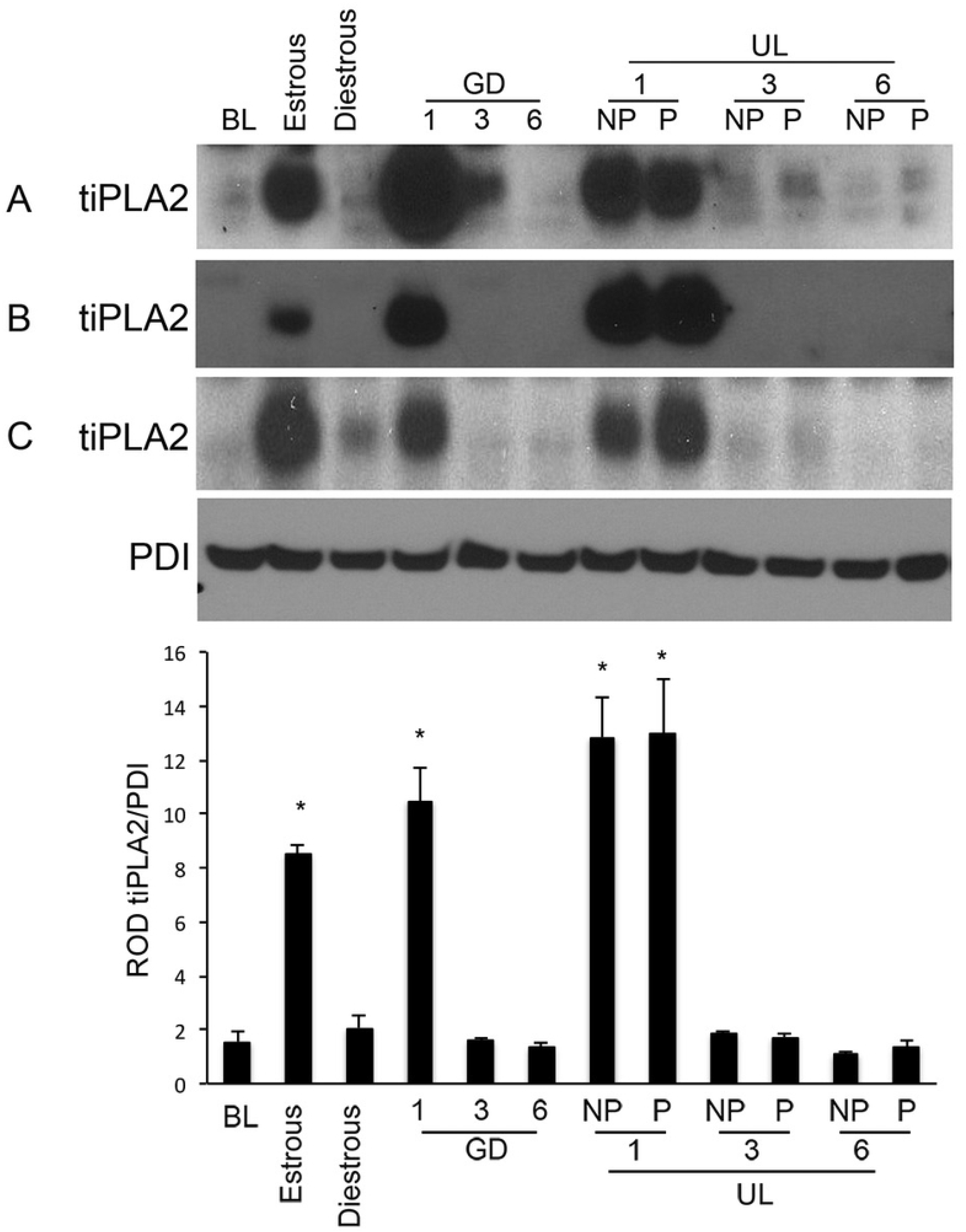
Uterine cleaved phospholipase A2 (tiPLA2) in control, unilateral and bilateral pregnant mouse models. Western blot analysis for tiPLA2 found at 20kda in mouse uterine tissue collected from bilateral ligated (BL) at 1 dpc, estrous, diestrous, and control pregnant mice at 1, 3, 6 dpc and unilateral ligated (UL), gravid and non gravid horns (P and NP respectively) on 1, 3, 6 dpc (n=3 for each gestational time point). Protein disulfide isomerase (PDI) is utilized as a cytoplasmic protein loading control. Representative blots from this experiment are shown. Statistical comparisons were performed using one-way ANOVA, and subsequent Newman–Keuls multiple-comparison tests. Data labeled with * are significantly elevated in comparison to BL uterine horn samples (*p* < 0.005).

### Uterine caspase-3 activation is both endometrial and myometrial in early gestation

Immunohistochemical analysis examined the presence of cleaved active caspase-3, in mouse uterine tissues isolated from our bilateral ligated mouse model (BL) 1 dpc, at estrous, diestrous and our unilateral ligated mouse model (UL) gravid (P) and non-gravid (NP) uterine horns at 1, 3 and 6 dpc (Figure 4). As can be observed caspase-3 activation on 1 dpc in the UL mouse model is strikingly isolated to the endometrial compartment in both the P and NP uterine horns as indicated by the pink/red staining of endometrial epithelial cells. At each other time point examined the endometrial epithelial cells remained free of caspase-3 activity. Isolated cells demonstrating caspase-3 activity could also be found in some glandular structures and endometrial stroma in all tissues examined (supplemental figures 1 and 2) and active caspase-3 is also found in the myometrial compartment increasing in intensity on 6dpc in the gravid (P) uterine horn. Non-ligated control mice (GD) 1, 3 and 6 demonstrated an identical pattern of caspase-3 activation as the UL (P) model (data not shown).

**Figure 4.**
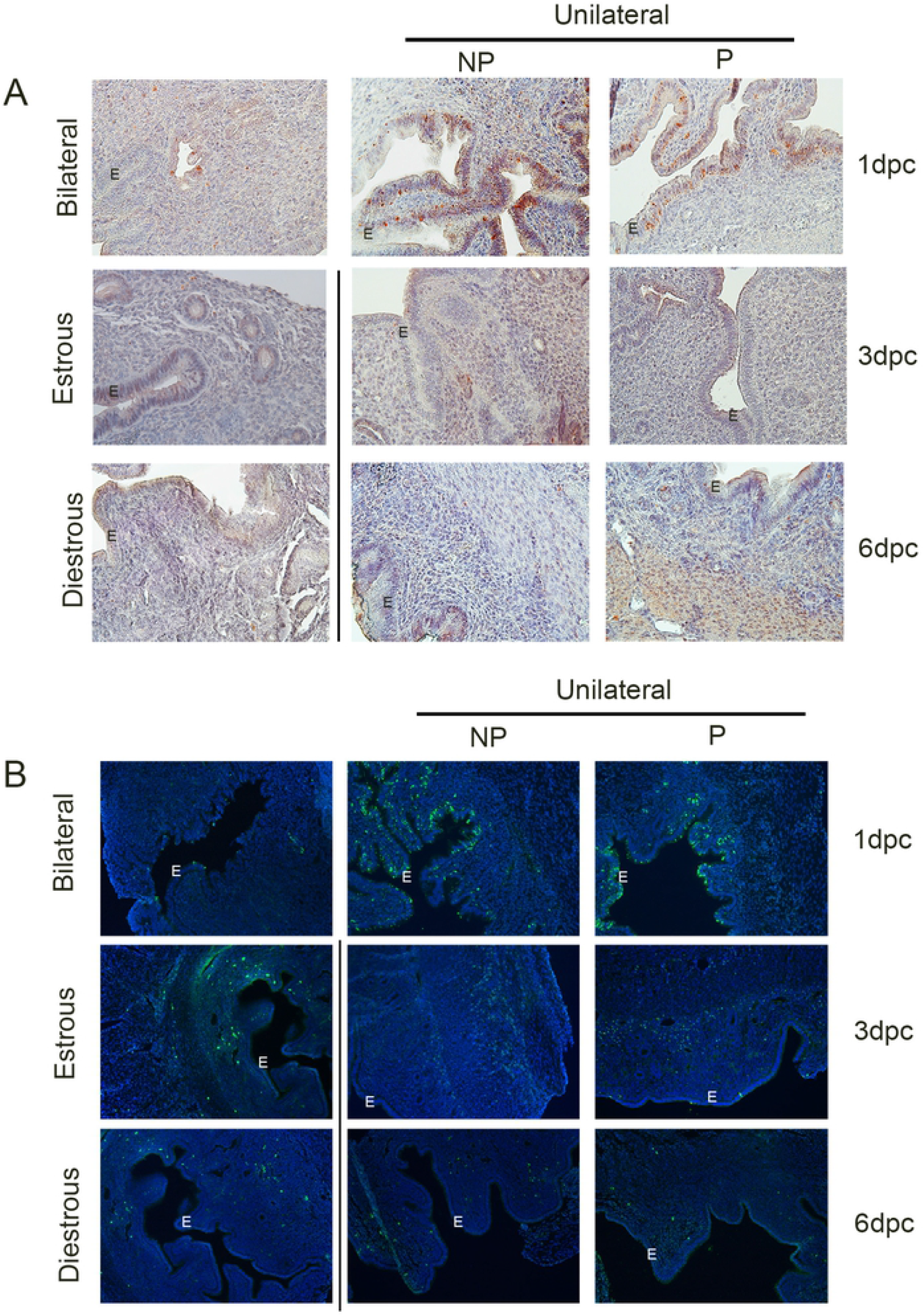
**A. Uterine cleaved caspase-3 immunohistochemical analysis in control, unilateral and bilateral pregnant mouse models.** Uteri isolated from estrous (positive control), diestrous (negative control), bilateral ligated (BL) 1dpc and unilateral ligated mouse models (UL), gravid and non-gravid (P/NP) horns on 1, 3 and 6 dpc, were examined by immunohistochemical analysis for cleaved caspase-3. A representative image is shown, magnification 20X. **B. Uterine apoptotic cell TUNEL analysis in control, unilateral and bilateral pregnant mouse models.** Mouse uteri isolated from estrous (positive control), diestrous (negative control), bilaterally ligated (BL) 1dpc and unilateral ligated mouse model (UL) gravid and non-gravid (P/NP) horns on 1, 3 and 6 dpc, were examined by TUNEL analysis for the presence of apoptotic cells. A representative image is shown, magnification 20X.

### Apoptosis is isolated to the endometrial compartment in early gestation

TUNEL staining (green) was performed to identify the site of apoptotic caspase-3 activity in mouse uterine tissues isolated from our bilateral ligated mouse model (BL) 1 dpc, at estrous, diestrous and our unilateral ligated mouse model (UL) gravid (P) and non-gravid (NP) uterine horns at 1, 3 and 6 dpc (Figure 4B). Nuclei were counterstained with Dapi (Blue). As can be observed apoptotic caspase-3 activation as indicated by TUNEL staining on 1dpc in the UL mouse model is isolated to the endometrial compartment in both the P and NP uterine horns as indicated by the fluorescent green staining of endometrial epithelial cells and some glandular structures. Though limited numbers of apoptotic cells were observed with equal frequency isolated to the endometrial stroma, in all other timepoints examined the endometrial epithelial cells and myometrial compartment remained free of apoptotic caspase-3 activity. Non-ligated control mice (GD) 1, 3 and 6 demonstrated the same pattern of TUNEL staining (data not shown). The lack of endometrial TUNEL staining in the BL uterine horn also confirms that coitus does not trigger apoptotic consequences in the endometrial compartment at least on 1dpc. Non-ligated control mice (GD) 1, 3 and 6 demonstrated an identical pattern of caspase-3 activation as the UL (P) model (data not shown).

## Discussion

This current study was initiated by our observation of a significant pre-implanation surge in active caspase-3 levels on 1 dpc in the pregnant mouse model (Figure 1), which rapidly declines to barely detectable levels on 2 dpc, prior to a mid gestational resurgence (5-17 dpc). As caspase-3 is the final effector enzyme in the apoptotic signaling cascade, which when activated can lead to programed cell death [11], we examined the consequence of its uterine activity in early gestation. Our previous analysis has determined the mid gestational surge (5-17dpc) in uterine caspase-3 activity is isolated to the myometrial compartment, is non-apoptotic and plays a critical tocolytic role, maintaining uterine quiescence across gestation [4, 5]. In contrast, at term and postpartum, uterine caspase-3 activity is apoptotic [12] in nature and limited to the endometrial compartment where it triggers a surge in prostaglandin biosynthesis through the truncation and activation of iPLA2 [6]. Importantly our previous analysis also determined that apoptotic caspase-3 possesses the ability via a proteolytic process to truncate and activate iPLA2 (amino acids 514-806), which consequently acts to generate lysophosphatidic acid and arachidonic acid from membranous phospholipids [13, 14], whereas non-apoptotic caspase-3 was unable to cleave and activate iPLA2. Cyclooxygenases (COX) 1 and 2 then convert the released arachidonic acid into PGH2, the common intermediate metabolite for biosynthesis of various prostaglandins [15].

Figure 2 confirms that similar to term and postpartum uterine caspase-3, uterine pre-implantation caspase-3 activity is also apoptotic in nature, (as indicated by elevated levels of PARP cleavage) in both the uterine horns isolated from control (GD) and the unilateral ligated (UL) pregnant model (P and NP) on day 1 dpc and at estrous (positive control). However, the absence of apoptotic caspase-3 activity in the bilaterally ligated model on 1 dpc demonstrates that the trigger for apoptotic caspase-3 activity is not coitus or the presence of sperm but suggests it is possibly triggered by a secreted circulating signal as a result of the presence of a conceptus. As we have previously identified uterine prostaglandin biosynthesis as a target of uterine apoptotic caspase-3 action [6] we asked the question as to whether pre-implantation apoptotic uterine caspase-3 performed a similar function. Indeed we found from prior observations by others that an early gestation pre-implanation surge in prostaglandin biosynthesis occurs with a PGE2 predominance and acts in a leutoprotective manner in early pregnancy by binding EP2 receptors on the corpus luteum increasing LH receptors and thereby sustaining increased progesterone secretion in early pregnancy and enhancing uterine receptivity [1, 16–18]

However, until now the molecular pathway that acts as a trigger for early uterine prostaglandin synthesis prior to implantation was unknown. Our analysis demonstrates that the pre-implanation surge in uterine apoptotic caspase-3 activity triggered by the presence of a fertilized gamete at 1dpc may play a central role in the increased levels of leutoprotective prostaglandin biosynthesis through its initial apoptotic targeting of uterine iPLA2.

By western blot analysis we determined the relative levels of uterine iPLA2 activation by examining for the presence of truncated iPLA2 (tiPLA2). As can be observed in figure 3, tiPLA2 was robustly upregulated only in the presence of apoptotic caspase-3 (Figure 3) i.e., in both the uterine horns isolated from control (GD) and the unilateral ligated pregnant model (P and NP) on day 1 dpc and at estrous (positive control).

Immunohistochemical analysis (Figure 4A) demonstrates that preimplantation caspase-3 activity is largely isolated to the endometrial compartment as indicated by the abundant dark red staining observed in the epithelial cell lining of the endometrial compartment both the unilateral ligated pregnant model (P and NP) on 1 dpc. Individual caspase-3 positive cells were also found in the stroma of the endometrium at each gestational time point. Widespread cytoplasmic myometrial caspase-3 was found at each gestational time point, which increased in intensity from 1-6 dpc in the gravid and decreased in intensity from day 1-6 in the non-gravid uterine horn of the unilaterally pregnant model (supplemental figure 1), recapitulating the levels of uterine active caspase-3 detected by western blot analysis in Figure 2. This likely represents non-apoptotic caspase-3 activity as confirmed by Figure 4B.

TUNEL staining (Figure 4B) confirms that the apoptotic activity in the preimplantation uterus is largely restricted to the endometrial compartment, as intense green TUNEL staining largely recapitulates the endometrial caspase-3 staining pattern observed in the epithelial cell lining of the endometrial compartment both the unilateral ligated pregnant model (P and NP) (Figure 4A). Individual TUNEL positive cells were also found in the stroma of the endometrium at each gestational time point. However, TUNEL staining was absent from the myometrial compartment. Previous studies have demonstrated a role for caspase-3 action in supporting a successful implanation event in that an intraluminal application of caspase-3 inhibitor inhibits implantation both in the hamster and mouse [19]. Thus, it appears that caspase-3-mediated uterine epithelial cell apoptosis plays an important role during the time of establishment of implantation. Moreover in support of the role of early uterine prostaglandins as mediators of implantation several human studies also demonstrate that using non-steroidal anti-inflammatory drugs (NSAIDs) which inhibit prostaglandin synthesis around the time of conception, significantly increases the risk of miscarriage by more than 4 fold in the first eight weeks of pregnancy [20]. Similarly in the pregnant mouse and rat models, administration of prostaglandin inhibitors block successful implantation events [21].

Other studies have demonstrated that inhibition of prostaglandin synthesis or receptor activation impairs endometrial implantation and causes early pregnancy loss. Peluffo et. al demonstrated that a treatment of female macaques with a selective PGE2 receptor antagonist had a contraceptive effect with a significantly reduction in rate of pregnancy compared to the control [22].

This work has important implications in understanding and postulating potential molecular mechanisms underlying impaired uterine receptivity observed in patients with implantation failure and early pregnancy loss. We recognize that early pregnancy involves a complex dynamic interplay between the embryo and endometrium with timing and compartment specific expression of factors requisite factors involved in endometrial receptivity. It is also well known that sufficient and properly timed increase in progesterone synthesis by the corpus luteum is necessary for endometrial decidualization and receptivity. However, the trigger for such upregulation of corpus luteum function remained elusive until now. Taken together our data demonstrates for the first time that a surge in endometrial apoptotic caspase-3 acts to initiate pre-implantation leutoprotective uterine prostaglandin biosynthesis in very early pregnancy. We propose that future studies are warranted to define cohorts of women who demonstrate a failure to activate endometrial caspase-3 apoptotic action as a risk factor for a failure to implant.

---

**Supplemental Figure 1. Uterine cleaved caspase-3 immunohistochemical analysis in mouse.** Immunohistochemistry staining in mouse uteri isolated from estrous (positive control), diestrous (negative control), bilaterally ligated (BL) 1 dpc and unilaterally ligated mouse model (UL) gravid and non-gravid (P/NP) horns on 1, 3 and 6 dpc. Magnification 10X

**Supplemental Figure 2. Stromal and myometrial mouse uterine cleaved caspase-3 immunohistochemical analysis.** Immunohistochemical analysis of mouse uteri isolated from estrous (positive control), diestrous (negative control), bilaterally ligated (BL) and unilaterally ligated (UL) mouse model gravid and non-gravid (P/NP) horns on 1, 3 and 6 dpc. Magnification, 20X

